# Modelling the role of dual specificity phosphatases in Herceptin resistant breast cancer cell lines

**DOI:** 10.1101/528315

**Authors:** Petronela Buiga, Ari Elson, Lydia Tabernero, Jean-Marc Schwartz

## Abstract

**Background:** Breast cancer remains the most lethal type of cancer for women. A significant proportion of breast cancer cases are characterised by overexpression of the human epidermal growth factor receptor 2 protein (HER2). These cancers are commonly treated by Herceptin (Trastuzumab), but resistance to drug treatment frequently develops in tumour cells. Dual-specificity phosphatases (DUSPs) are thought to play a role in the mechanism of resistance, since some of them were reported to be overexpressed in tumours resistant to Herceptin.

**Results:** We used a systems biology approach to investigate how DUSP overexpression could favour cell proliferation and to predict how this mechanism could be reversed by targeted inhibition of selected DUSPs. We measured the expression of 20 DUSP genes in two breast cancer cell lines following long-term (6 months) exposure to Herceptin, after confirming that these cells had become resistant to the drug. We constructed several Boolean models including specific substrates of each DUSP, and showed that our models correctly account for resistance when overexpressed DUSPs were kept activated. We then simulated inhibition of both individual and combinations of DUSPs, and determined conditions under which the resistance could be reversed.

**Conclusions:** These results show how a combination of experimental analysis and modelling help to understand cell survival mechanisms in breast cancer tumours, and crucially enable us to generate testable predictions potentially leading to new treatments of resistant tumours.

## Background

Breast cancer is one of the most common and the most lethal type of cancer affecting women. Approximately one in eight women in the Western world develops breast cancer throughout her life [1]. HER2-positive cases represent about 25%, characterised by high levels of HER2 activity arising from mutations, overexpression of the HER2 protein or amplification of the *HER2* gene [2, 3].

HER2 (Neu or ErbB2) belongs to the protein tyrosine kinase (PTK) epidermal growth factor receptor family that consists of three other proteins: HER1, HER3 and HER4 [3]. All four HER receptors function as homo-or hetero-dimers that are activated by a variety of ligands, such as the epidermal growth factor (EGF), transforming growth factor a (TGFα), heparin-binding EGF-like growth factor and neuregulins (NGFs) [4, 5]. HER2 incorporates into heterodimers with the other HERs resulting in its activation. No ligand of HER2 has been discovered yet [6]. HER2 can also be activated by ligand-independent homodimerization that follows its overexpression in tumour cells [7]. Active HER2 auto-phosphorylates and binds to various molecules to activate signalling pathways, for example the Mitogen-Activated Protein Kinase (MAPK) and phosphatidylinositol 3-kinase (PI3K) pathways, which collectively promote cell growth, proliferation and survival [8].

Because of their critical roles in driving cancer, PTKs are major targets for therapy that is often administered in the form of small molecule inhibitors or neutralizing antibodies [9, 10]. HER2-positive breast cancer is commonly treated by Herceptin (Trastuzumab), a humanized monoclonal antibody that associates with the extracellular domain IV of the protein. Dimerization of HER2 is decreased by Herceptin, thereby inhibiting PI3K and MAPK pathways, promoting antibody-mediated cellular cytotoxicity, and promoting HER2 ubiquitinylation and internalisation [8]. One third of HER2-positive breast cancer cases are responsive to Herceptin, but two thirds of these relapse within one year due to resistance to the drug that develops in the tumour cells [11, 12] [13]. Potential mechanisms for resistance to Herceptin include overexpression of other tyrosine kinases that replace HER2, structural alterations of HER2 that remove or mask the Herceptin binding-site on HER2, or alterations in downstream signalling pathways that reduce their dependence on HER2 [8-10]. Many efforts to overcome this resistance have been investigated including combination of Herceptin with other drugs that target HER2 (e.g. Lapatinib) or key downstream proteins (e.g. inhibitors of Raf, MEK and PI3K) [9, 14].

In tumour cells, HER2 signalling is regulated by several MAPKs, most importantly ERK1, ERK2, p38, and JNK. These kinases are activated by phosphorylation of specific threonine and tyrosine residues by upstream kinases, and inactivated by dephosphorylation of either or both residues. Dephosphorylation is carried out by the dual-specificity phosphatases (DUSPs) which belong to the tyrosine phosphatase superfamily [15]. The DUSP family is comprised of ten MAPK phosphatases (MKPs) and of additional atypical DUSPs, which actively down-regulate MAPK activity [16, 17] [18]. Individual DUSPs have distinct subcellular localization and substrate specificity. Many DUSPs were linked with various cancer types [19] and some of them, such as DUSP4, have been reported to be overexpressed in tumours resistant to Herceptin [20].

In order to study Herceptin resistance in breast cancer we turned to two HER2-positive human breast cancer cell lines: BT-474, which is estrogen receptor (ER) and progesterone receptor (PR) positive, and SK-BR-3, which is ER and PR negative [21, 22]. These differences determine unique clinical outcomes, specific responses to therapeutic strategies and specific progression of metastasis [23]. Reports indicate that exposing these cells to Herceptin leads to development of resistance to the drug after periods that range from 3 to 12 months [24-26] [27, 28] [20, 29]. The molecular mechanisms that lead to resistance in these cell lines, including the possible involvement of DUSPs in this process, are unknown.

Computational modelling is a well-recognised approach to study regulation of cell signalling processes and to predict their effects [30, 31] [32]. Creating kinetic models of signalling pathways is often challenging because the detailed dynamics and interactions of these pathways are usually not known, and complex experimentation is required to provide the missing data. An efficient alternative is to use Boolean models [33], which reproduce the dynamics of the system in a qualitative manner by discretising levels of nodes into two possible states: 0 or 1. Interactions are similarly discretised into activation or inhibition [33]. We have previously constructed a series of Boolean models for short-term exposure of cells to Herceptin, which correctly accounted for the described regulatory mechanisms involving DUSPs [34]. The present study uses Boolean models combined with experimental data to identify DUSPs that may be targeted in breast cancer cells for reversing Herceptin resistance.

## Methods

### Cell lines

The human breast cancer cell lines BT-474 and SK-BR-3 were purchased from the American Type Culture Collection (Manassas, USA). BT-474 cells were cultured in DMEM medium containing 50 U/ml penicillin, 50 mg/ml streptomycin and 10% heat-inactivated foetal bovine serum (FBS – Gibco/ThermoFisher Scientific, Waltham, USA). SK-BR-3 cells were cultured in McCoy’s 5A medium (Sigma-Aldrich, St. Louis, USA), supplemented as above. Both cell types were grown at 37°C in an atmosphere of 5% CO2.

### Generation of Herceptin resistant cell lines

Cells were grown in duplicate in 60 mm plates and cultured for 48 hours until they were 75% confluent. Following that, they were exposed to 50 µM Herceptin (Roche, Switzerland) on an ongoing basis for six months. Cells were fed every 72 hours with Herceptin-containing medium and split when confluency reached 75%. Cells grown in parallel for the six-month period in the absence of Herceptin were used as controls.

### Proliferation assay of Herceptin sensitivity

Aliquots of cell cultures that had been grown for six months in the presence or absence of Herceptin were plated in 6-well plates. Following overnight adherence, a count of viable cells was taken. The remaining wells of each culture were then grown for 72 hours in the presence and absence, respectively, of 50 µM Herceptin, after which cells were counted once again. The change in cell numbers in the presence and absence, respectively, of Herceptin was calculated.

### Gene expression analysis by q-PCR

Total cellular RNA was prepared using the RNeasy Mini Kit (QIAGEN, Germany); RNA was treated with DNaseI prior to use. One microgram of RNA was reverse transcribed using the qScript cDNA synthesis kit (Quanta Biosciences, Beverly, USA) in a total volume of 20 µl according to the manufacturer’s instructions. Quantitative PCR (RT-qPCR) experiments were carried out using 0.5 µl (25 ng) cDNA with the KAPA SYBR Fast qPCR Master Mix ABI Prism (Kapa Biosystems, Wilmington, USA) together with target-specific primers (as described in [34]). Amplification was performed on an AB StepOnePlus instrument (Applied Biosystems, Foster City, USA). The amplification conditions consisted of an initial activation step of 95°C (20 seconds), followed by 40 x (95°C, 3 seconds; 60°C, 30 seconds) and ending with 1 x (95°C, 15 seconds; 60°C, 60 seconds; 95°C, 15 seconds).

ΔCT (average change in threshold cycle number) values were determined for each DUSP in each sample relative to endogenous controls (β-actin ad GAPDH) by the ΔΔCT method [35]. Experiments were performed in triplicate using two biological repeats. The Primer3 software was used to design DUSP-specific forward and reverse primers [36, 37], and their efficiency was assessed by standard curves. Student’s t-test was used to determine significance by the GraphPad Prism software (GraphPad Software, La Jolla, USA).

### Construction of Boolean models

Boolean models were manually constructed and run as described previously [34]. Cell survival was represented by adding a specific Survival node in the model. We added an activation link between ERK and Survival, since in healthy cells it is observed that ERK, JNK and p38 favour proliferation. In addition, we added an interaction that represents combined interaction of JNK and p38 inhibiting Survival. This is justified because the high levels of activity of JNK and p38 favour apoptosis and inhibition of cellular growth [38-41]. When the Survival node is ON, the outcome should be interpreted as a set of cellular processes favouring proliferation; when the Survival node is OFF, the outcome should be interpreted as a set of cellular processes favouring cell death.

When both JNK and p38 are inhibited by DUSPs, cell death is prevented in tumour cells. We used information about DUSPs and their specific substrates published in the literature to connect them with their known kinases in our models [18, 42]. In order to simulate an overexpressed DUSP we introduced an unknown activator “A”, which was kept ON during the whole time course. To simulate the inhibition of a targeted DUSP we introduced a node “I”, which had an inhibitory action on its target and was kept ON permanently.

## Results

### Induction of Herceptin resistance by long-term treatment with the drug

In order to determine if long-term treatment with Herceptin induces resistance to the drug in the BT-474 and SK-BR-3 cell lines, cells from each line were cultured continuously for six months in the absence and presence, respectively, of 50 µM Herceptin. Massive cell death was not observed during this prolonged period in cells treated with Herceptin. Then, aliquots of both treated and non-treated cells were exposed to 50 µM Herceptin for 72 hours, and their proliferation during this period was quantified. In agreement with previous reports [20, 24-28, 30], Herceptin inhibited proliferation of BT-474 and SK-BR-3 cells that had not been exposed to long-term Herceptin treatment. In contrast, proliferation of cells that had been exposed for 6 months to Herceptin was not affected by this additional 72-hour treatment with Herceptin (Figure 1), confirming that they are resistant to the effects of the drug.

**Figure 1.**
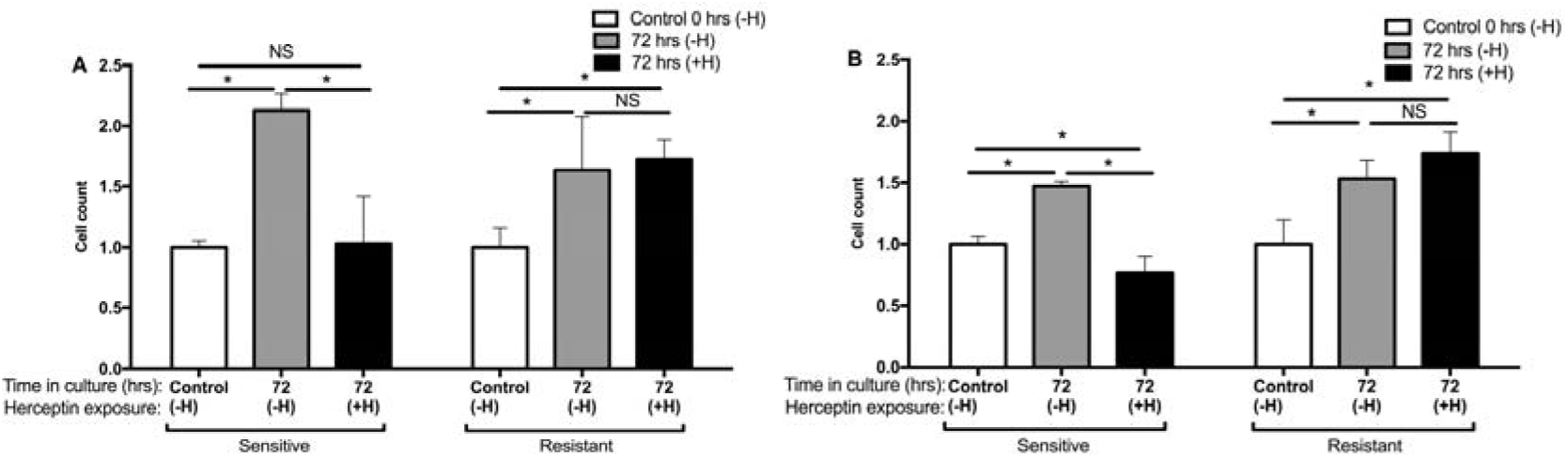
Herceptin sensitivity of resistant BT-474 (A) and SK-BR-3 (B) cell lines. 24 hours after seeding in the absence of Herceptin, cell aliquots were counted (Control). Other aliquots were grown for an additional 72 hours with (+H) or without (-H) 50 µM Herceptin and then counted. Cells had previously been grown in culture for six months in the absence and presence, respectively, of 50 µM Herceptin (marked as Sensitive or Resistant, respectively). Bars represent mean ± SE cell numbers, relative to Control, of two experiments each performed in triplicate. * means p < 0.05. The growth of sensitive, but not resistant, BT-474 and SK-BR-3 cell lines was inhibited by Herceptin.

### Resistance to Herceptin alters *DUSP* expression in BT-474 cells

Resistance to Herceptin may arise following changes in signalling downstream to HER2. In order to determine if Herceptin resistance alters expression levels of DUSPs, we quantified the expression of 20 *DUSP* genes (10 *MAPK* phosphatases and 10 atypical *DUSP*s) using RT-qPCR in Herceptin-resistant and Herceptin-sensitive BT-474 and SK-BR-3 cell lines. Within each cell line, expression of a given *DUSP* was quantified in resistant cells relative to its expression in the sensitive cells. Data for BT-474 cells are presented in Figure 2.

**Figure 2.**
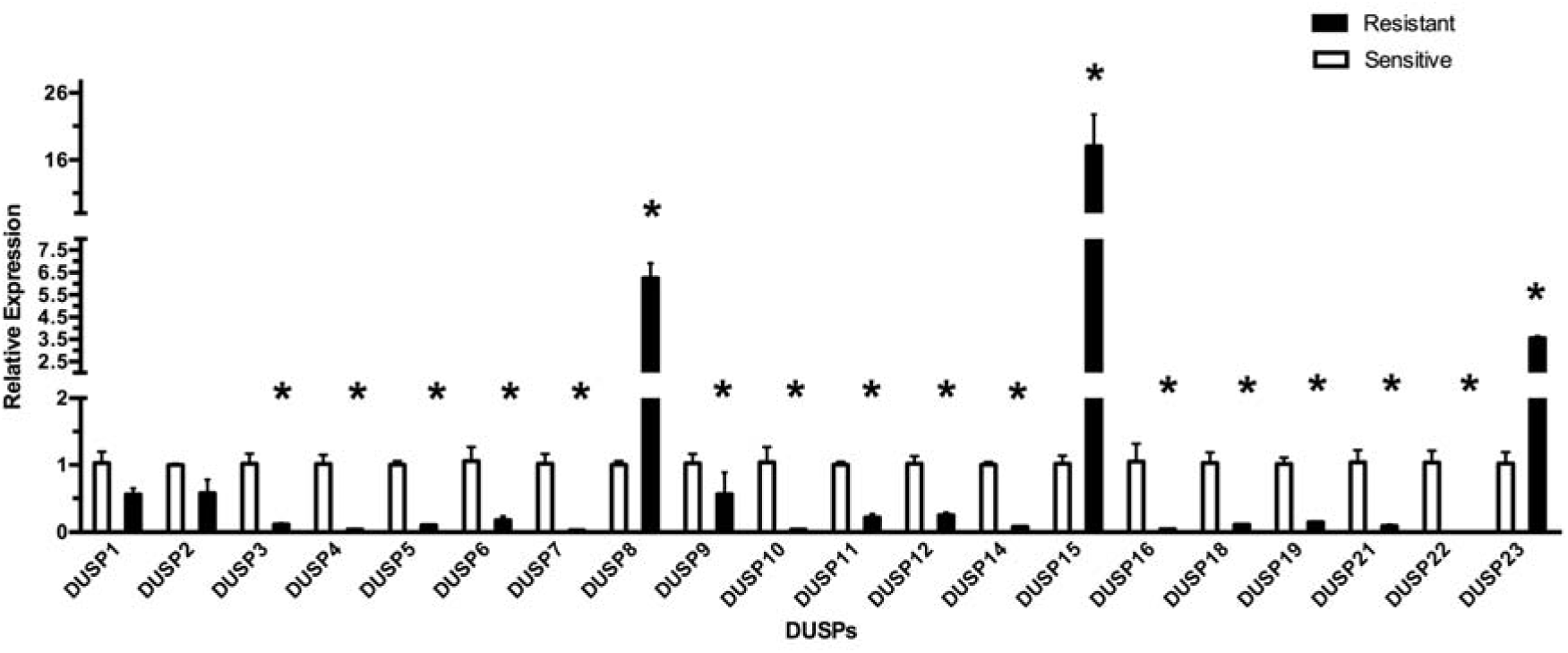
*DUSP* expression in BT-474 cells. Shown is the expression of each *DUSP* measured by RT-qPCR in cells rendered Herceptin-resistant by treatment with the drug for six months, normalized to its expression in Herceptin-sensitive cells. Bars indicate mean ± standard error (SE). * means p < 0.05 between treated and non-treated cells.

Following Herceptin treatment, we observed fluctuating levels of individual *DUSP* expression and a significant reduction in expression of a large number of *DUSPs* in the BT-474 cell line, including *DUSP*s 3, 4, 5, 6, 7, 9, 10, 11, 12, 14, 16, 18, 19, 21 and 22. However, expression of *DUSPs* 8, 15 and 23 was significantly increased.

### Modelling selective inhibition of DUSPs in BT-474 cells

Boolean models were constructed to represent and simulate resistance to Herceptin *in vitro*. As indicated in Methods, increased expression of a *DUSP* in Herceptin resistant cells was simulated by introducing an unknown activator (A). In a conceptually-similar manner, down-regulation of a *DUSP* was simulated by introducing an unknown inhibitor (I), as shown in Figure 3 (A-D).

**Figure 3.**
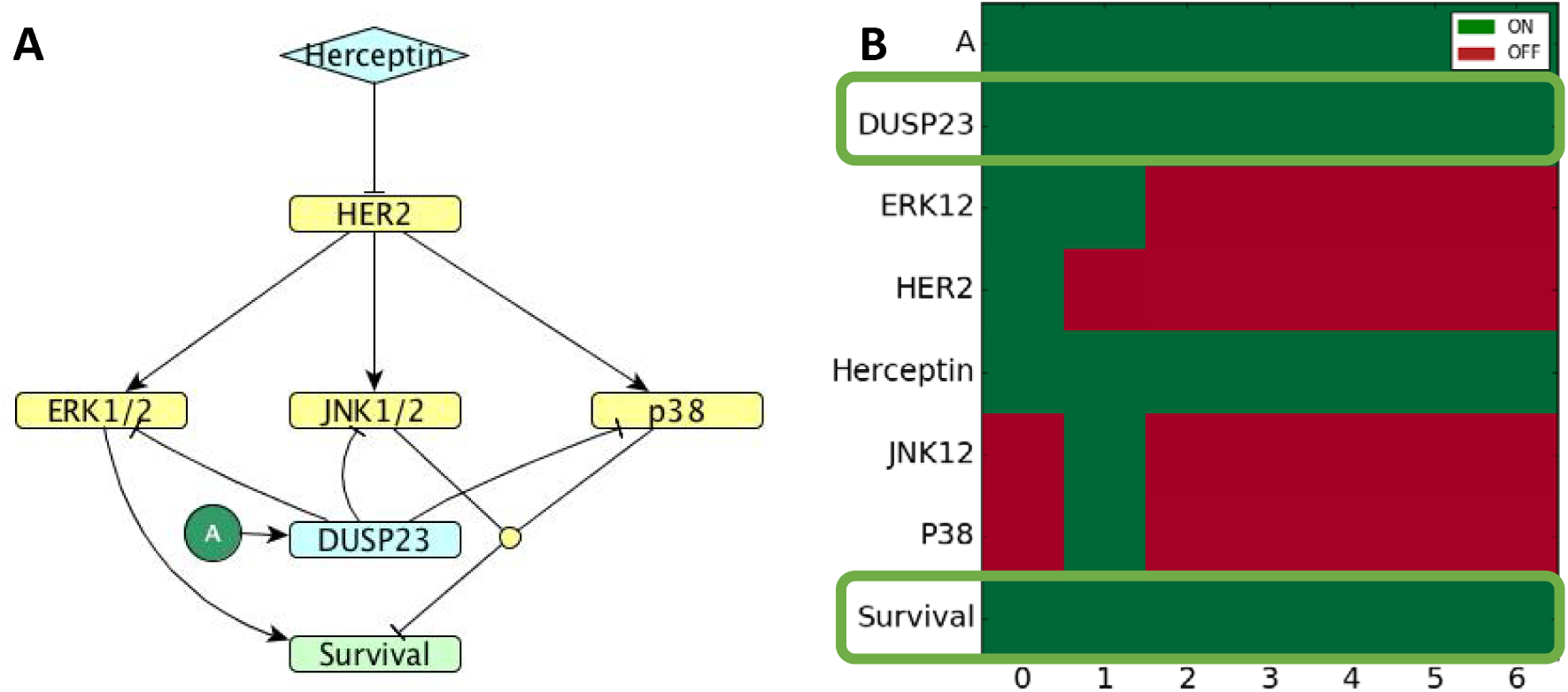
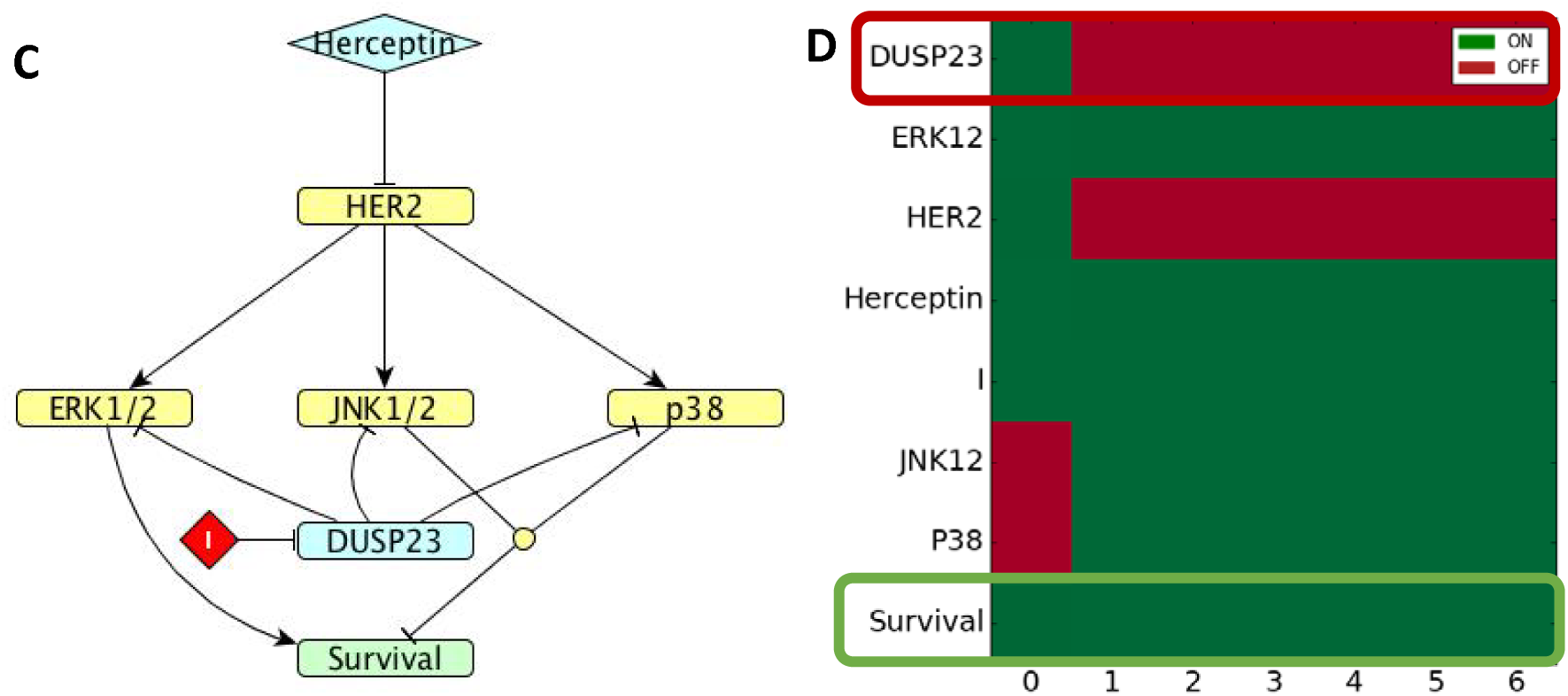
A. *DUSP*23 model simulating resistance. Graph of *DUSP*23 regulation in BT-474 cells treated with Herceptin. Overexpressed *DUSP*23 is represented by an inducer “A”. B. Gene expression simulation results where the state of each node is represented in green (ON) and red (OFF). C. Model of *DUSP*23 inhibition in resistance. Graph of *DUSP*23 regulation in BT-474 cells treated to Herceptin. “I” represent the inhibitor of *DUSP23*. D. Gene expression simulation results as in Figure 3.B. The horizontal axis represents arbitrary time units in all simulations.

We focused our studies on DUSPs that are up-regulated in the Herceptin-resistant state and whose inhibition might reverse resistance, in light of continuing efforts to design DUSP inhibitors [43, 44]. As a first step, we show that our models are consistent with the observed property of resistance. *DUSP23* substrates are known to be ERK [45], JNK1/2 and p38 [46], but the specific MAPK kinase involvement in *DUSP23* induction is not known. Our model shows that up-regulation of *DUSP23* expression is consistent with resistance to the drug by keeping Survival ON (Figure 3, A, B). However, inhibition of DUSP23 expression did not change this outcome, as cells are predicted to survive in this case as well (Figure 3, C, D). DUSP23 is therefore predicted not to be a suitable target for reversing Herceptin resistance.

### Identification of potential targets to reverse Herceptin resistance in BT-474 cells

In case of DUSP15, there are no known substrates from among the MAP kinases; this phosphatase regulates the ERK1/2 transduction pathway most likely via intermediary factors [47], hence we did not model this *DUSP*.

*DUSP8* inhibits JNK and p38 [42], but the specific MAPK kinase involvement in *DUSP8* induction is not known (Figure 4, A and B). DUSP8 is overexpressed in Herceptin-resistant BT-474 cells (Figure 2), and when applying an activator to the DUSP, we confirmed that Survival is maintained ON in our model. When applying an inhibitor to DUSP8, the model predicts that Survival is switched OFF, indicating that it is a possible target for reversing resistance (Figure 4, C and D). Moreover, this outcome remains unchanged if we introduce into the model both *DUSPs* 8 and 23 and inhibit only *DUSP8* (Figure 3, E and F).

**Figure 4.**
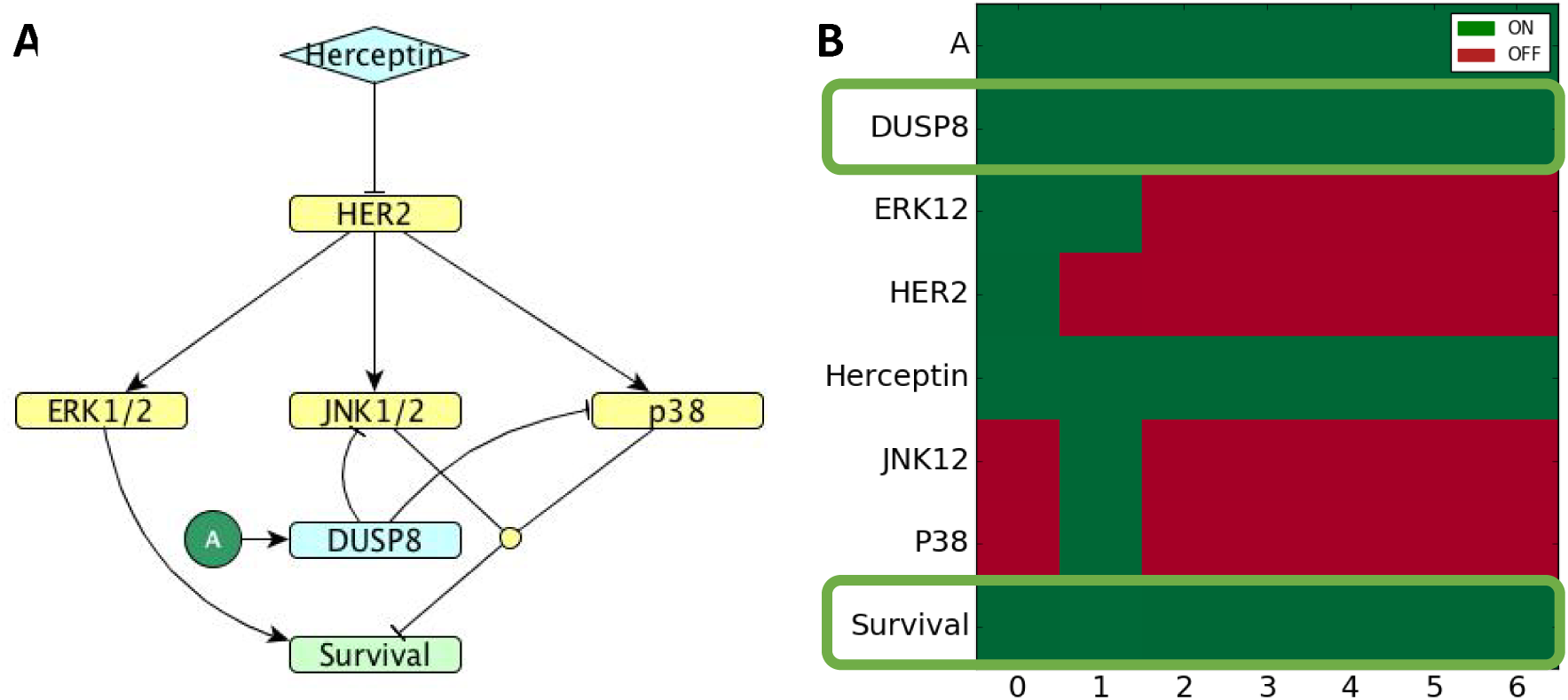
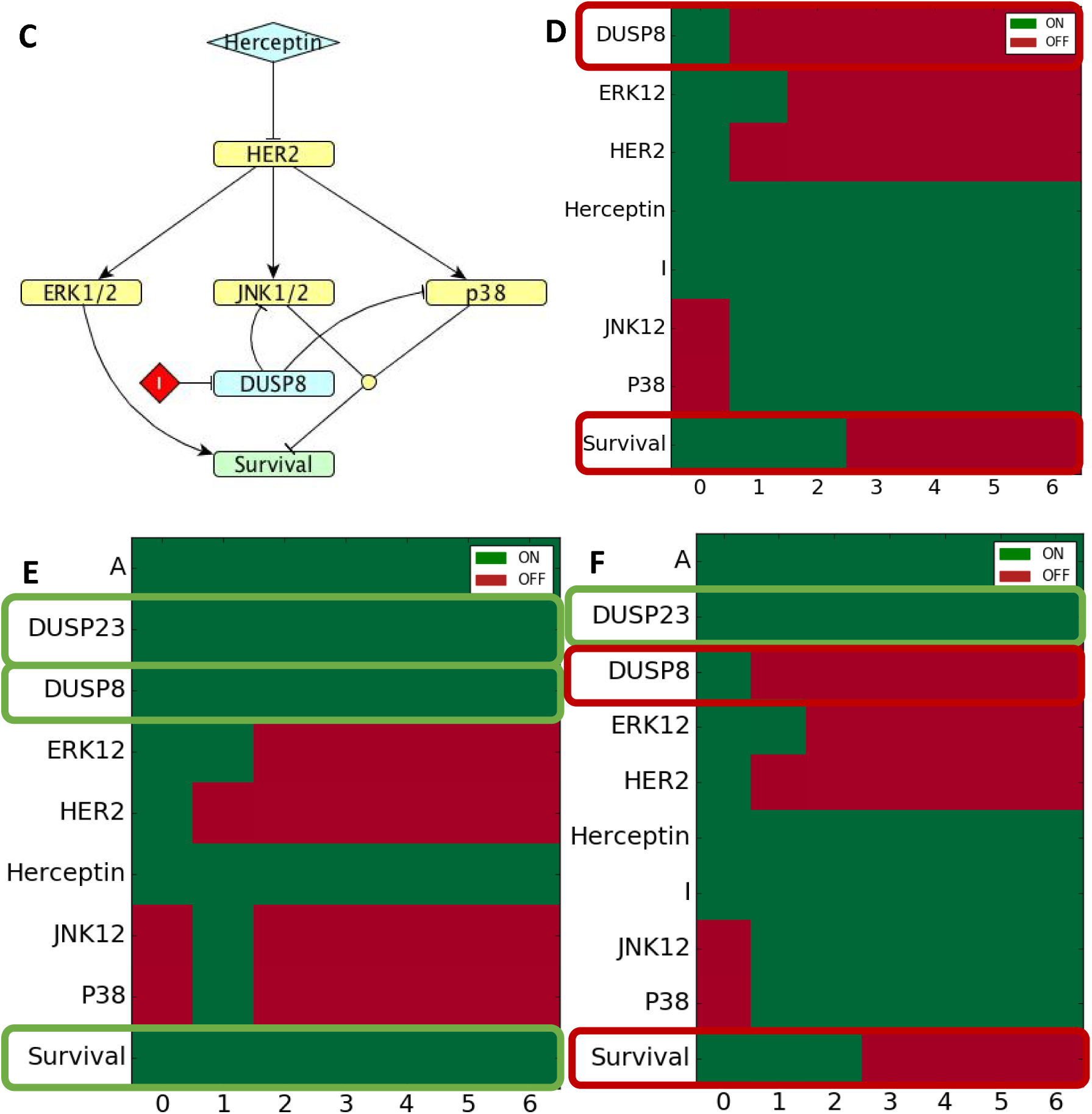
A. *DUSP8* model simulating resistance. Graph of *DUSP8* regulation in BT-474 cells exposed to Herceptin. Overexpressed *DUSP8* is represented by an inducer “A”. B. Gene expression simulation results as in Figure 3.B. C. Model of *DUSP8* inhibition in resistance. Graph of *DUSP*8 regulation in BT-474 cells exposed to Herceptin. “I” represents inhibition of *DUSP8*. D. Gene expression simulation results as in Figure 3.B. E. Resistance model, *DUSP*8 and 23 are overexpressed and Survival is ON. F. Model of *DUSP*8 inhibition in resistance, *DUSP*23 is not inhibited and Survival is OFF. The horizontal axis represents arbitrary time units in all simulations.

### *DUSP* expression is altered by Herceptin resistance in SK-BR-3 cells

Similar to the BT-474 cell line, expression levels of the 20 *DUSP*s determined by RT-qPCR in the SK-BR-3 cell line are shown in Figure 5.

**Figure 5.**
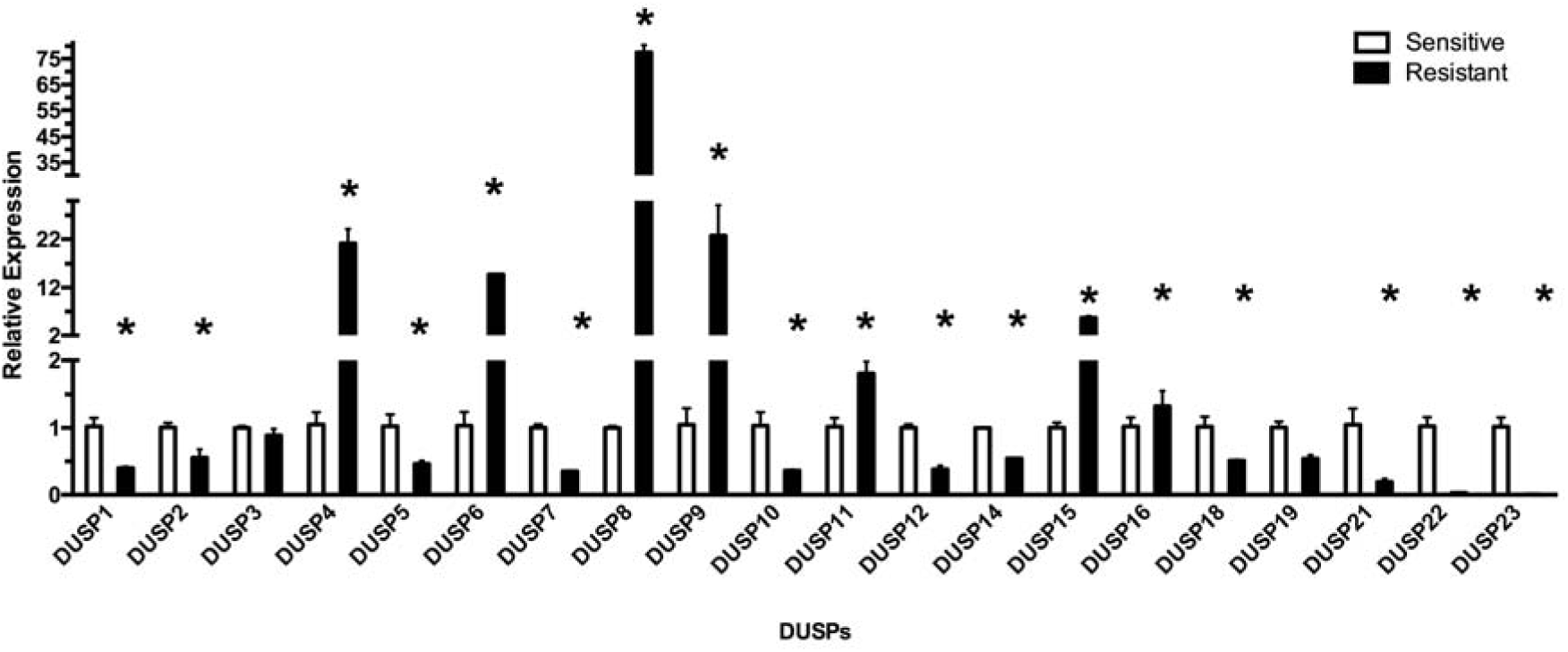
*DUSP* expression in SK-BR-3 cells. Shown is the expression of each *DUSP* measured by RT-qPCR in cells rendered Herceptin-resistant by treatment with the drug for six months, normalized to its expression in Herceptin-sensitive cells. Bars indicate mean ± standard error (SE). * means p < 0.05 between treated and non-treated cells.

Resistance to Herceptin increased expression of more DUSPs in SK-BR-3 cells compared to BT-474 cells. Expression of *DUSPs* 4, 6, 8, 9, 11, 15 and 16 was significantly increased; of note, this list includes DUSPs 8 and 15, which were also up-regulated in Herceptin-resistant BT-474 cells. Expression of *DUSPs* 1, 2, 5, 7, 10, 12, 14, 18, 22 and 23 was significantly decreased.

### Modelling selective inhibition of DUSPs in SK-BR-3 cells

We simulated overexpression of DUSPs for which regulatory mechanisms were known by applying an activator “A” and tested if inhibition of these DUSPs could reverse survival. We modelled the role of overexpressed DUSPs 4 and 6 in resistance separately in SK-BR-3 cells (Figure 6, A-H). *DUSP4* and *DUSP6* are induced by ERK1/2 [42]. *DUSP4* inhibits ERK1/2 and JNK1/2, while *DUSP6* inhibits only ERK1/2 [42]. Our simulations indicate that resistance is not overcome by inhibiting these DUSPs separately, as cell survival is maintained.

**Figure 6.**
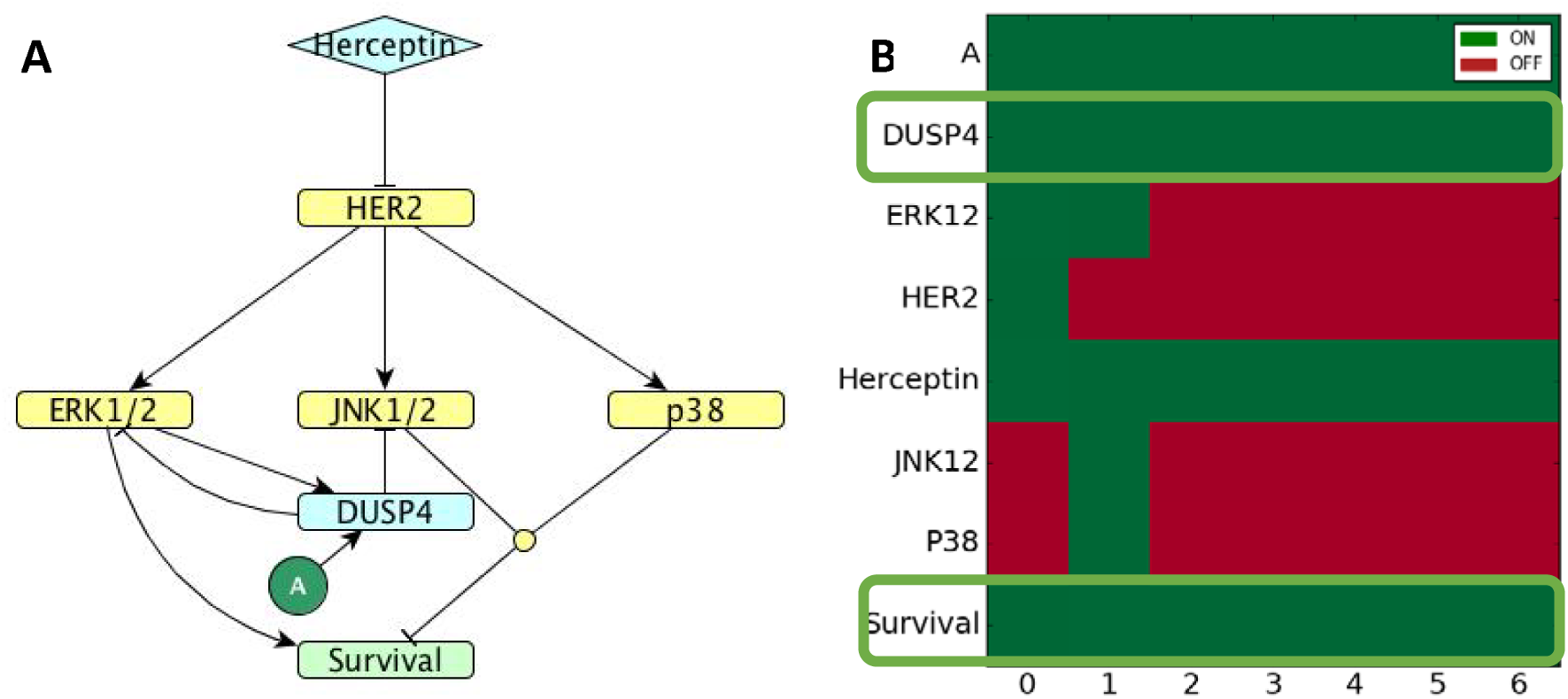
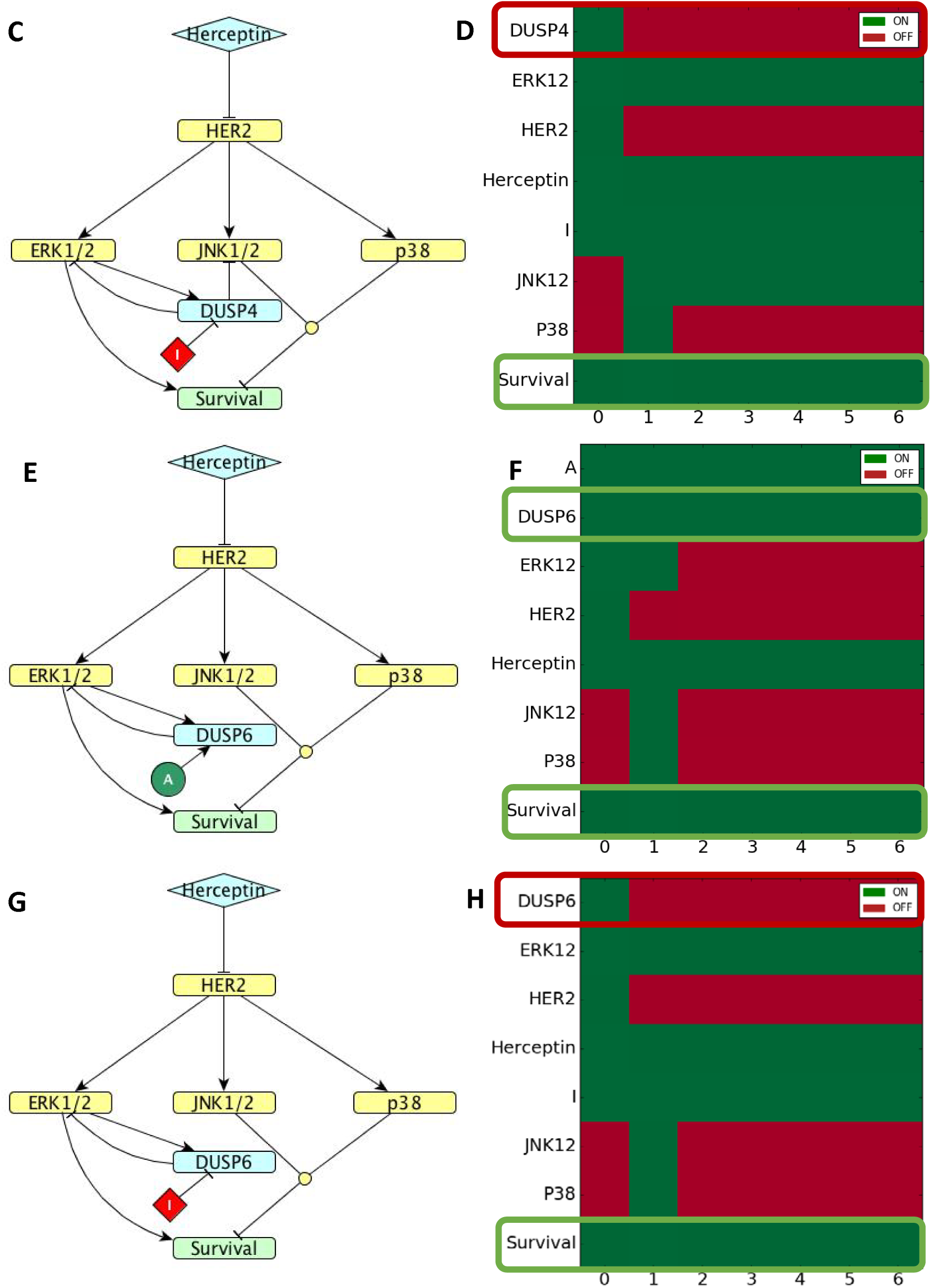
A. *DUSP4* model simulating resistance. Graph of *DUSP4* regulation in SK-BR-3 cells exposed to Herceptin. Overexpressed *DUSP4* is represented by an inducer “A”. B. Gene expression simulation results as in Figure 3.B. C. Model of *DUSP4* inhibition in resistance. Graph of *DUSP*4 regulation in SK-BR-3 cells exposed to Herceptin. “I” represents inhibition of *DUSP4*. D. Gene expression simulation results as in Figure 3.B. E-H. Similar to A-D, respectively, for *DUSP6*.

Simulating the effects of overexpression and inhibition of DUSP8 indicated that, as was observed in BT-474 cells, Survival remains “ON” when this DUSP is overexpressed, but changes to “OFF” when this DUSP is inhibited (Figure 7, A, B). Similar effects were noted also for overexpression and inhibition of DUSP16 (Figure 7, C, D). *DUSP8* and *DUSP16* inhibit JNK and p38 [42], but the specific kinase involvement in their induction is not known. We further confirmed that irrespective of the activation of other DUSPs, such as DUSPs 4 and 6, Survival is switched OFF if DUSP8 is inhibited (Figure 7, C, D).

**Figure 7.**
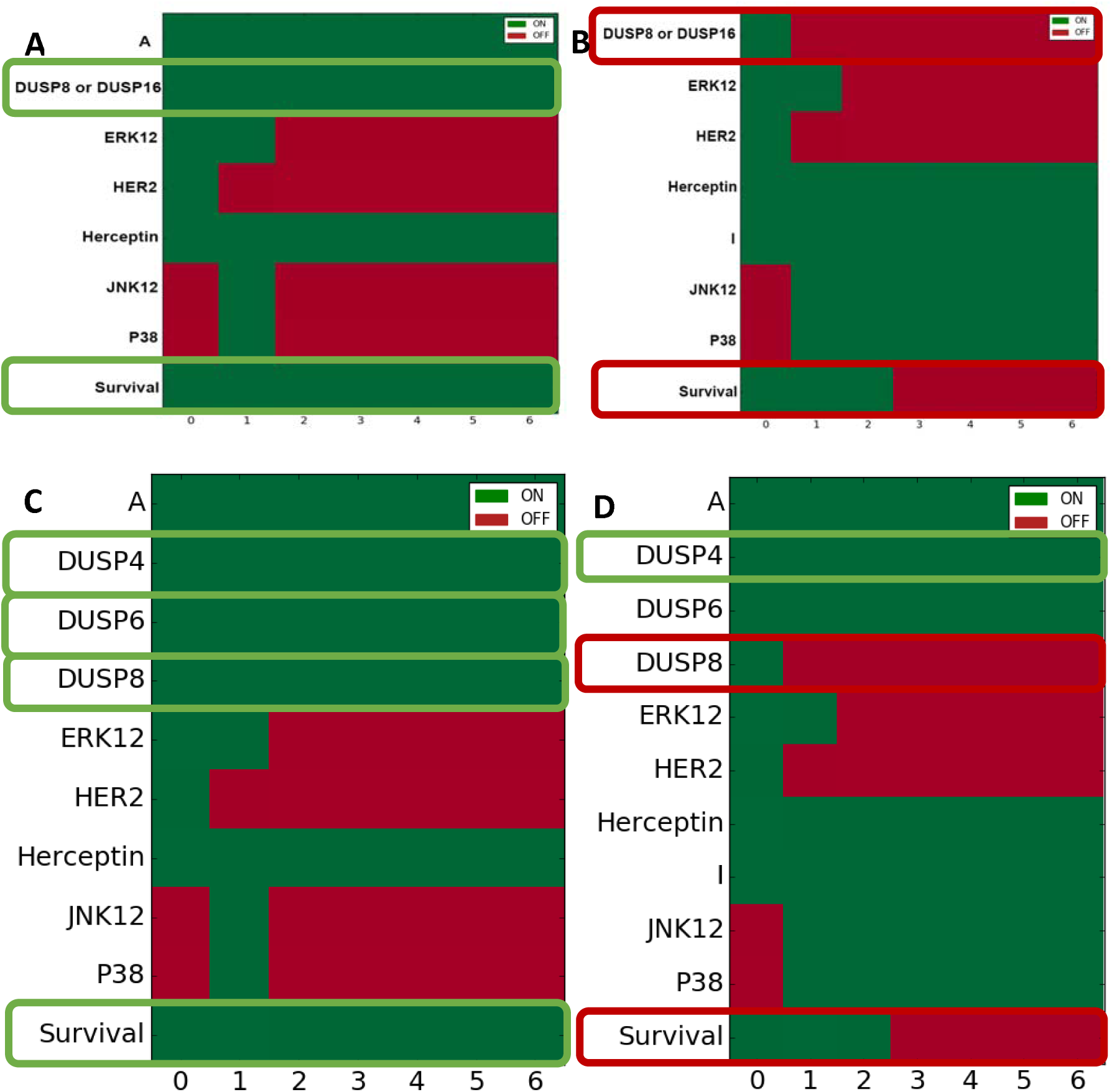
Simulation of the Herceptin-resistance model of SK-BR-3 cells. A. *DUSP8* or *DUSP16* are overexpressed and Survival is ON. B. Inhibition of either *DUSP8* or *DUSP16* (note addition of “I” row in heatmap) changes Survival to OFF. C.

Overexpression of *DUSPs 4, 6* and *8* retains Survival as ON. D. Modelling inhibition of *DUSP8* in the presence of overexpressed *DUSP4* and *DUSP6* changes Survival to OFF. The horizontal axis represents arbitrary time units in all simulations.

## Discussion

In a previous study, we built several Boolean models to study the initial response of HER2-positive breast cancer cells to Herceptin and the contribution of DUSPs in pathways that signal cell survival [34]. We were able to explain the observed dynamics in the expression of several DUSPs playing a role in regulation of MAPK signalling and to predict new regulatory mechanisms for other DUSPs. In the present study we use a similar approach to examine the roles of DUSPs in Herceptin resistance in the BT-474 and SK-BR-3 human breast cancer cells that had been rendered Herceptin resistant by long-term treatment with the drug. We observed that expression of most *DUSPs* was downregulated upon long-term Herceptin exposure. However, several *DUSPs* were significantly overexpressed in one or both cell types, including *DUSPs* 4, 6, 8, 9, 11, 15, 16 and 23. Among these, *DUSP*s 4, 6, 8, 9 and 16 are classical MKPs that target MAPK kinases and therefore we expect that they may function in controlling cell survival. The rest are atypical DUSPs, whose role in cell survival is less well known.

We constructed several Boolean models that incorporated established regulatory mechanisms of *DUSPs.* We confirmed that they were able to simulate the property of resistance to Herceptin. The models also enabled us to predict inhibition of which of the *DUSPs* overexpressed in Herceptin-resistant cells would reverse resistance and render the cells once again sensitive to Herceptin. Our results indicate that inhibition of *DUSP8*, alone or in combination with other *DUSPs*, reverses Herceptin resistance. Inhibition of *DUSP16* produced similar results. The finding that *DUSP8* was overexpressed in two disparate breast cancer cell lines further strengthens its relevance to this issue. Both DUSP8 and DUSP16 share the substrates JNK1/2 and p38, therefore both are able to switch the Survival node to OFF when inhibited, separately or in combination with other DUSPs. The other *DUSPs* we found to be overexpressed did not share the same combination and specificity of substrates. Another MKP with similar substrates is DUSP10, but its expression was not upregulated in resistant cells. It is worth noting that we simulated other combinations of DUSP upregulation, but the conclusions of our models remained unchanged if other DUSPs were upregulated, as long as either DUSP8 or DUSP16 was upregulated.

DUSPs have been correlated with resistance to Herceptin by Györffy and colleagues, which highlighted DUSP4 as being involved in Herceptin resistance in HER2 positive breast cancer [48]. Menyhart et al. found high expression of DUSP4 and 6 to be correlated with worse survival of HER2-positive breast cancer patients. In that case, treatment with Herceptin and transiently silencing *DUSP4* simultaneously induced sensitivity to Herceptin in resistant cell lines [20].

Our results enable us to propose a possible strategy to avoid development of Herceptin resistance, if suitable inhibitors for these DUSPs can be found. The effect of DUSP silencing on cellular behaviour must be further investigated experimentally. These results show how a systems biology approach can lead to better understanding of mechanisms responsible for cell proliferation in breast cancer tumours and generate testable hypotheses potentially leading to new treatments of resistant tumours.

## Authors’ contributions

PB conducted experiments, analysed data, developed models and wrote the manuscript. JMS, AE and LT conceived the study and edited the manuscript. All authors read and approved the final manuscript.

## Competing interests

The authors declare no competing interests.

## Acknowledgements

PB is funded by Weizmann UK and by University of Manchester. We acknowledge funding from a “Making-Connections” grant from Weizmann-UK (P17231) and a grant from the Lord Alliance “Get Connected” programme between the University of Manchester and the Weizmann Institute. AE is incumbent of the Marshall and Renette Ezralow Professorial Chair.

